# Diverse Early Life-History Strategies in Migratory Amazonian Catfish: Implications for Conservation and Management

**DOI:** 10.1101/018671

**Authors:** Jens C. Hegg, Tommaso Giarrizzo, Brian P. Kennedy

## Abstract

Animal migrations provide important ecological functions and can allow for increased biodiversity through habitat and niche diversification. However, aquatic migrations in general, and those of the world’s largest fish in particular, are imperiled worldwide and are often poorly understood. Several species of large Amazonian catfish carry out some of the longest freshwater fish migrations in the world, travelling from the Amazon River estuary to the Andes foothills. These species are important apex predators in the main stem rivers of the Amazon Basin and make up the regions largest fishery. They are also the only species to utilize the entire Amazon Basin to complete their life cycle. Studies indicate both that the fisheries may be declining due to overfishing, and that the proposed and completed dams in their upstream range threaten spawning migrations. Despite this, surprisingly little is known about the details of these species’ migrations, or their life history. Otolith microchemistry has been an effective method for quantifying and reconstructing fish migrations worldwide across multiple spatial scales and may provide a powerful tool to understand the movements of Amazonian migratory catfish. Our objective was to describe the migratory behaviors of the three most populous and commercially important migratory catfish species, Dourada (*Brachyplatystoma rousseauxii*), Piramutaba (*Brachyplatystoma vaillantii*), and Piraíba (*Brachyplatystoma filamentosum*). We collected fish from the mouth of the Amazon River and the Central Amazon and used strontium isotope signatures (^87^Sr/^86^Sr) recorded in their otoliths to determine the location of early rearing and subsequent. Fish location was determined through discriminant function classification, using water chemistry data from the literature as a training set. Where water chemistry data was unavailable, we successfully in predicted ^87^Sr/^86^Sr isotope values using a regression-based approach that related the geology of the upstream watershed to the Sr isotope ratio. Our results provide the first reported otolith microchemical reconstruction of *Brachyplatystoma* migratory movements in the Amazon Basin. Our results indicate that juveniles exhibit diverse rearing strategies, rearing in both upstream and estuary environments. This contrasts with the prevailing understanding that juveniles rear in the estuary before migrating upstream; however it is supported by some fisheries data that has indicated the presence of alternate spawning and rearing life-histories. The presence of alternate juvenile rearing strategies may have important implications for conservation and management of the fisheries in the region.

## Introduction

Animal migration provides many important ecological functions: they can be a stabilizing strategy in seasonal environments; offer transitory habitats for large populations; often transport materials across ecosystem boundaries; and may increase a regions biodiversity [1]. Large-scale migrations shed light on ecosystem connectivity across scales and can be used as a lens to understand broader behavioral responses to the environment and links to physical processes [2–4]. However, migrations worldwide are under threat from the alteration of migratory pathways, habitat loss, climatic changes and anthropogenic changes to the landscape [5]. In aquatic systems, changes in upstream land use and the placement of dams have had significant impacts on ecosystems and migrations worldwide [6–11]. This is particularly true for large migratory fish that are under threat in many of the world’s largest river systems [12–14]. Despite this, many large migratory fish species are not well understood [14].

Globally, dams and water resources challenges in the two largest rivers in China provide an example of the ongoing changes to large rivers and their effects on aquatic species, including sturgeon and paddlefish [15]. In South America, transnational river systems and a lack of coordinated research of aquatic systems may result in losses to unspecified levels of biodiversity [16–20]. New dams present a unique challenge to migratory fish in the region. Because the young of many Amazonian species undergo a drifting larval stage, even if adults can pass above dams the lack of flow in reservoirs creates a barrier that drifting juveniles are unable to surmount on their way downstream [21].

Several species of Amazonian catfish in the genus *Brachyplatystoma* carry out some of the longest freshwater fish migrations in the world, travelling over 4,500 km from rearing areas in the Amazon estuary to spawning grounds in rivers in the foothills of the Andes [22–24]. These species largely inhabit whitewater and clearwater rivers within the Amazon Basin [sensu 25,26], with rare reports in tannic blackwater rivers [23]. These catfish species are the only known organisms, terrestrial or aquatic, that require the entire length of the Amazon basin to complete their life cycle [23]. They are also one of the few apex predators in the pelagic and demersal zones of the largest Amazonian rivers, playing an important role in trophic dynamics and ecosystem functioning within the entire basin [27]. However, evidence indicates that the fisheries for the most populous species are in decline, potentially due to overfishing [22,28]. The reliance of these species on headwater streams for spawning leaves adults and larva vulnerable to blocking of their migration paths by dams and their reservoirs [21,29].

Surprisingly little is known about the life history of migratory Amazonian catfish given that the three most abundant *Brachyplatystoma* taxa support the largest fisheries in the Amazon Basin [23,24]. Dourada (*Brachyplatystoma rousseauxii*) is a pelagic predator found throughout the whitewater and clearwater rivers of the Amazon and supports the largest fishery in the Amazon [27,29]. Piramutaba (*Brachyplatystoma vaillantii*) make up a second large export fishery and are found almost exclusively in the Amazon River mainstem, whitewater tributaries, and the estuary [30–32]. Piraíba (*Brachyplatystoma filamentosum*) is the largest of the migratory catfish, present in whitewater rivers throughout the Amazon basin. It is also the most locally exploited and least understood of these three species [22]. The expansive scale of the Amazon Basin, and the large size of the rivers these fish inhabit, have made tracking and reconstructing their movements very difficult [32]. Our current understanding of the migratory behavior of migratory Amazonian catfish is based on fishing records (including the catch timing and size of fish across the Amazon basin) and a growing number of scientific sampling efforts [23,28,31,33–36]. After hatching in the upper reaches of the whitewater rivers originating in the Andes, larvae of these species drift downstream for two to four weeks before reaching the Amazon estuary. Juveniles rear in the estuary before commencing an upstream migration that coincides with the seasonal flood pulse. Genetic data indicate that dourada may home to natal tributaries in the basin to spawn [37,38].

Otolith microchemistry has been an effective method for quantifying and reconstructing fish migrations worldwide across multiple spatial scales [39–46]. Strontium ratio in particular has become a powerful tool for determining movement and location because it is not fractionated biologically. Thus, the signatures recorded in otoliths match the water through which fish pass [43,47–50]. Studies of geological weathering throughout the Amazon basin have provided detailed, multi-year records of micro-chemical and isotopic chemistry in the largest rivers of the basin. These data provide the required background sampling necessary to tie regional otolith signatures to geographic location [51–53] (Figure 1A, Table 1). Recent studies have also shown the feasibility of predicting ^87^Sr/^86^Sr signatures of unknown watersheds using the geologic makeup of the basin, allowing researchers to characterize strontium signatures of unsampled areas [54,55]. These advances point to otolith microchemistry as a potentially powerful tool to understand the movements of Amazonian migratory catfish.

**Figure 1.**
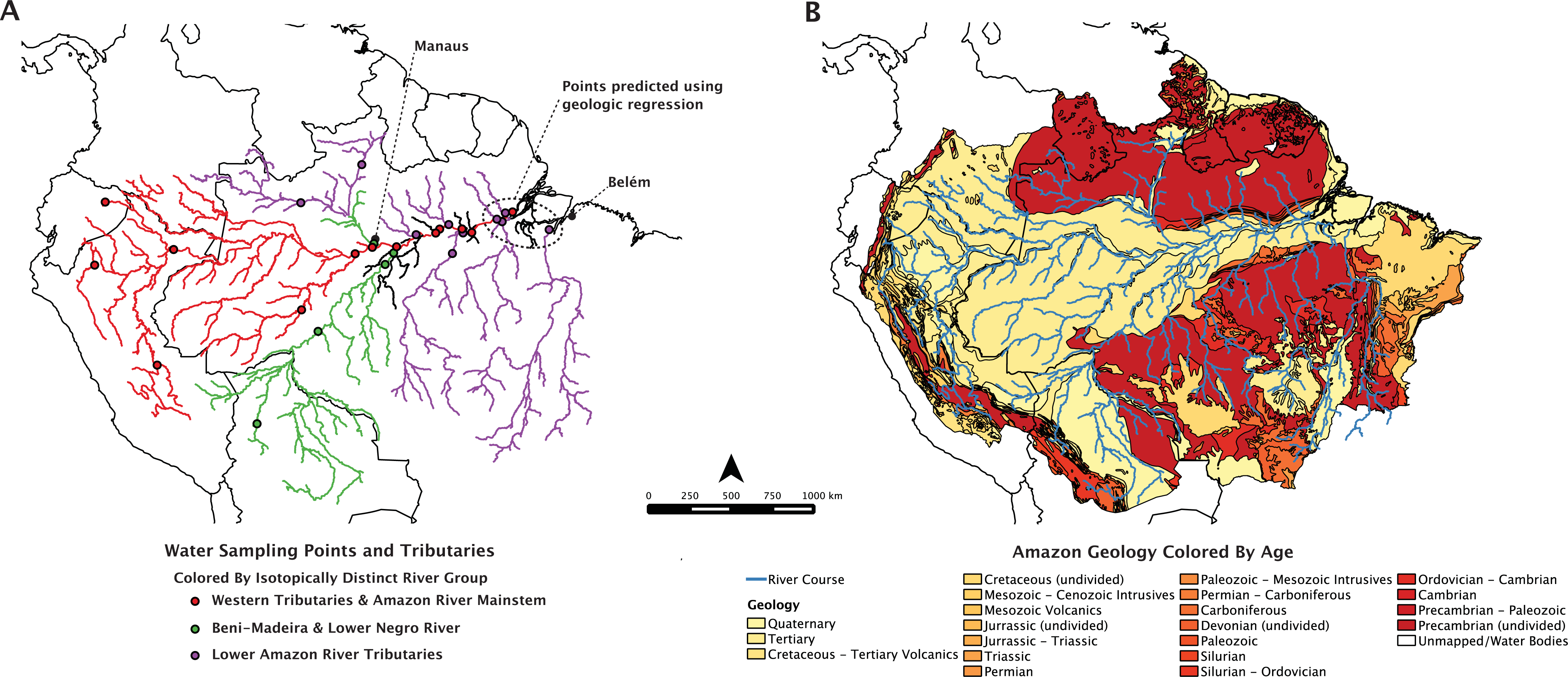
Water sampling points and geology of the Amazon River basin. Maps show (A) the location of ^87^Sr/^86^Sr water samples within the Amazon River basin digitized by the authors from location descriptions in the literature [51–53] and points predicted from Equation 1. The geological age and composition of the basin (B) used to predict the ^87^Sr/^86^Sr signatures of unsampled watersheds is also shown. Maps created using USGS datasets [59–61]

**Table 1.** *Brachyplaystoma* spp. sample information. Fish were collected from two locations in the Brazilian Amazon; in the cities of Belém near the mouth of the Amazon River, and Manaus in the central Amazon Basin. Piramutaba were gutted prior to otolith collection. Their weights are estimated from a length-to-weight relationship from Pirker [56].

| Species | Sample Number | Location | Total Length (cm) | Weight (kg) |
| --- | --- | --- | --- | --- |
| *Brachyplatystoma rousseauxii*<br>(*dourada*) | BR1 | Belém | 75 | 8.00 |
|  | BR3 | Belém | 75 | 8.00 |
|  | BR6 | Belém | 75 | 8.00 |
|  | BR7 | Belém | 75 | 8.00 |
|  | BR8 | Belém | 75 | 8.00 |
|  | BR10 | Belém | 75 | 8.00 |
|  | BR12 | Belém | 75 | 8.00 |
|  | BR14 | Belém | 75 | 8.00 |
|  | BR16 | Belém | 75 | 8.00 |
|  | BR18 | Belém | 100 | 22.00 |
|  | BR19 | Manaus | 110 | 11.00 |
|  | BR21 | Manaus | 100 | 6.00 |
|  | BR23 | Manaus | 115 | 10.00 |
|  | BR24 | Manaus | 75 | 3.70 |
|  | BR25 | Manaus | 80 | 4.00 |
|  | BR26* | Manaus | - | - |
| *Brachyplatystoma vaillanti*<br>(*piramutaba*) | BV1 | Manaus | 75 | 4.19° |
|  | BV2 | Manaus | 70 | 3.38° |
|  | BV3 | Manaus | 68 | 3.09° |
|  | BV4 | Manaus | 65 | 2.68° |
|  | BV5 | Manaus | 70 | 3.38° |
| *Brachyplatystoma filamentosum*<br>(*piraíba*) | BF1 | Belém | 220 | 110.00 |
|  | BF3 | Belém | 250 | 130.00 |
|  | BF5 | Belém | 90 | 20.00 |

\* Asterisci otolith. Analysis was excluded due to low Sr concentrations
° Weights are estimated from length-to-weight ratios

Our objective was to describe the migratory behaviors of large, migratory catfish in the Amazon River basin using otolith microchemistry. We focused our study on the three most populous and commercially important species in the Amazon Basin. We sought to determine the location of early rearing and subsequent movement in dourada, pirimutaba, and piraíba using samples collected from two, large fish markets at the mouth of the Amazon River and in the central Amazon. We determined the movement patterns over the lifetime of individual fish using laser ablation isotope mass spectrometry of their otoliths. Areas of stable signature were identified statistically throughout the chemical profile of the otolith, which were then classified to their location within the basin using discriminant function analysis. The discriminant function was created using a training set of ^87^Sr/^86^Sr samples from rivers throughout the Amazon basin. These samples were obtained from the geological literature. Where river ^87^Sr/^86^Sr values were unknown, we used established relationships between surface water ^87^Sr/^86^Sr values and the age and composition of the underlying watershed geology to predict these signatures.

## Ethical statement

Ethical approval was not required for this study, as all fish were collected as part of routine fishing procedures. Fish were sacrificed by the artisanal fishermen in Manaus and Belém using standard fisheries practices and donated to the authors.

No field permits were demanded to collect any samples from any location, since all samples derived from commercial catch. None of the species included in this investigation are currently protected or endangered. Therefore, no additional special permits were necessary. Permission to export the otolith samples was granted by the Brazilian Government with permit number: 116217 (MMA, IBAMA, CITES 09/01/2013).

## Methods

### Otolith Collection

In March 2012, a total of 24 paired lapillus otolith samples (16 pairs for dourada, 5 for piramutaba and 3 for piraíba) were collected from the two major fishing ports of Brazilian Amazonia, the cities of Manaus and Belém (Table 1, Figure 1A). These cities are located 1,606 river kilometers apart. Manaus (03°05′39.60″S, 60°01′33.63″W), is the largest city in the central Amazon, located at the confluence of the whitewater Solimões River with the blackwater Negro River. Belém (01°27′18.04″S 48°30′08.90″W), is situated on the banks of the Amazon estuary and is the main landing port of large migratory catfishes fisheries in Brazil [30]. Prior to otolith collection the total length (TL) and weight (W) of each fish was recorded: dourada: mean TL = 83.6 cm, mean W = 8.58 kg; piramutaba: mean TL = 67.6 cm, mean W = 3.34 kg; piraíba: mean TL =186.6 cm, mean W = 86.6 kg. Piramutaba were gutted prior to collection so weight was estimated using a length-to-weight ratio from Pirker [56].

Fish collected from Manaus were captured in the mainstem Amazon River between the mouth of the Madeira River and Manaus as reported by the fisherman. Thus, we would expect the chemical signatures representing the end of the fish’s life (signatures from the edge of the otolith) of fish caught in Manaus to represent signatures in the mainstem Amazon River or its tributaries upstream of the Madeira River. Fish collected in Belém were captured in the estuary between 60 km and 150 km from Belém according to the fisherman. The otolith edge chemistry of fish caught in Belém are therefore assumed to match the signature of the estuary, the lower Amazon tributaries, or the lower Amazon River mainstem. Because the ^87^Sr/^86^Sr signature can take days to weeks to equilibrate and accumulate enough material to reliably sample, it is possible that fish could exhibit signatures other than the location of capture if they had recently moved from a habitat with a different signature.

### Otolith Analysis

The left lapillus otolith from each sample was prepared using standard methods of mounting, transverse sectioning with a high precision saw, and abrasive polishing to reveal the rings [34,57] (Figure 2). If the left otolith was missing or unavailable the right otolith was used for analysis. Otoliths were then analyzed at the GeoAnalytical Laboratory at Washington State University using a Finnigan Neptune (ThermoScientific) multi-collector inductively coupled plasma mass spectrometer coupled with a New Wave UP-213 laser ablation sampling system (LA-MC-ICPMS). We used a marine shell standard to evaluate measurement error relative to the global marine signature of 0.70918 [58]. Repeated analyses of a marine shell signature provided an average ^87^Sr/^86^Sr value of 0.70914 during the course of the study (N=22, St. Error=0.00002). The laser was used to ablate a sampling transect from the core of the otolith section to the edge (30 μm/s scan speed, 40 μm spot size, 0.262 s integration speed, ∼7 J/cm). This resulted in a continuous time-series of ^87^Sr/^86^Sr data from the birth of the fish (core) to its death (edge) which was used for subsequent analysis. For more detailed methods see Hegg et. al [40]. The asteriscus was used for one sample for which the lapilli were not available; however, the strontium concentration was low and the unreliable results were not included.

**Figure 2.**
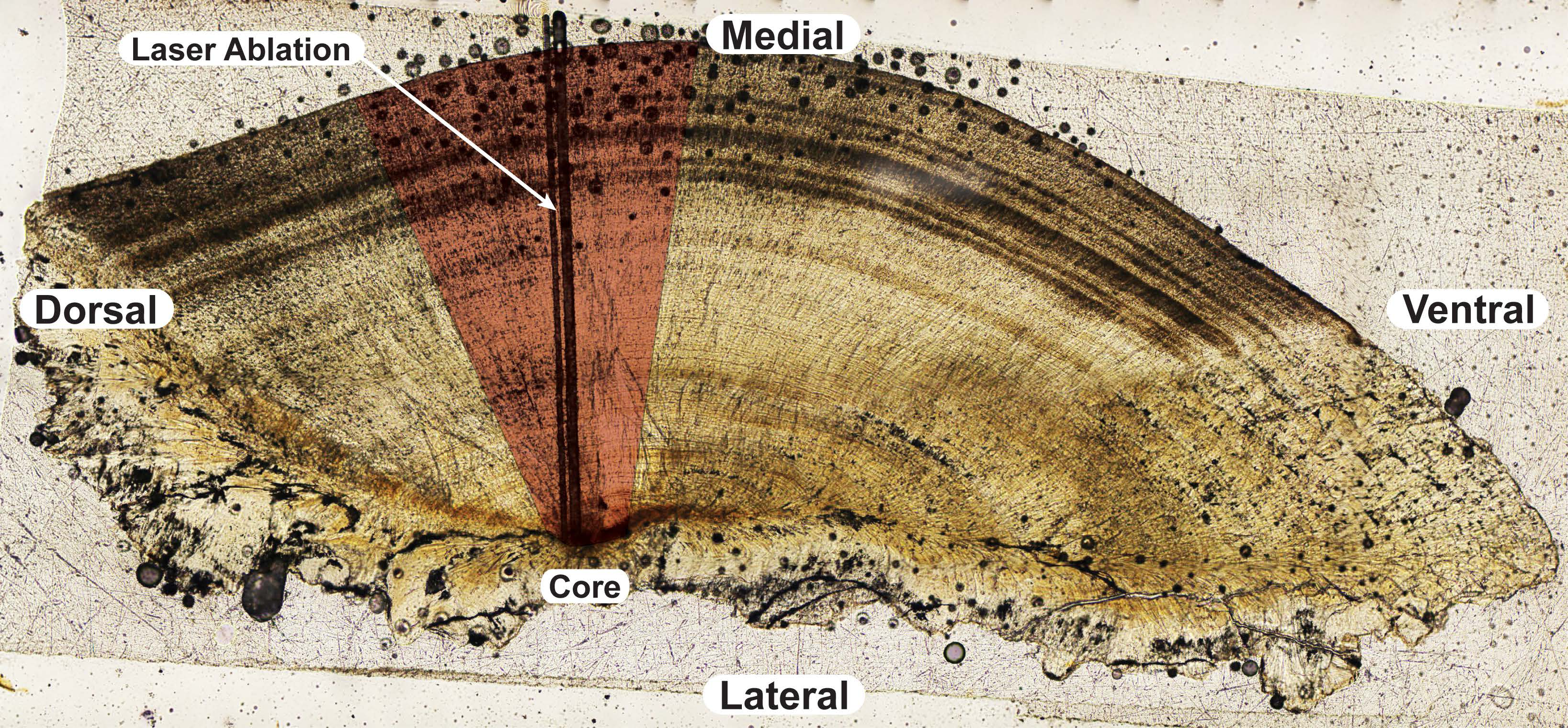
Otolith sectioning and analysis. Representative transverse section from a dourada lapillus otolith showing the analysis area (in red) used for all otoliths with the laser-ablation tracks indicated. All analyses were performed approximately perpendicular to the growth rings.

### Baseline Water Sampling and Prediction

Twenty-four water sampling points located throughout the Amazon River basin (Figure 1A) from three published studies provided baseline ^87^Sr/^86^Sr values for our study (Table 2, Figure 3). Santos et al. [53] provided thirteen samples from the Ore-HYBAM project (www.ore-hybam.org), a multi-year research effort with a comprehensive sampling design covering the mainstem Amazon River and all of the major tributaries above Obidos, Brazil. Nine additional samples from Gaillardet et al. [52] covered the mainstem Amazon and the mouths of the major tributaries as far east as Santarem, Brazil. Finally, Queiroz et al. [51] provided two samples from the Lower Solimões and Upper Purus Rivers. Our intention was to include samples that represented all major Amazon tributaries at a regional scale, while excluding smaller tributaries that were unlikely to provide long-term habitat for our study species. Smaller tributaries in the Amazon Basin have been shown to exhibit much different isotopic chemistry from their mainstem rivers [51]. The scale and geologic heterogeneity of these smaller tributaries could jeopardize assignment accuracy [54].

**Table 2.** Isotopic and geologic makeup of major watersheds of the Amazon River basin. Strontium ratios were taken from water samples reported in the literature for locations throughout the Amazon Basin and used as baselines to determine the likely location of fish movement. These samples were classified to three statistically distinguishable river group classifications using quadratic discriminant function analysis. Unsampled locations were predicted using geologic regression (Equation 1).

| Sample Point Name<br>(From Literature) | Literature Source | River | River Group Classification | $^{87}\text{Sr}/^{86}\text{Sr}$ | St. Dev. | N | Geology Percent Area | | | Mean Age<br>(Ma) |
| --- | --- | --- | --- | --- | --- | --- | --- | --- | --- | --- |
|  |  |  |  |  |  |  | Carboniferous | Tertiary | Precambrian |  |
| Amazon 13 | Gaillardet et al. 1997 | Amazon | Wester Tributaries & Amazon Mainstem | 0.710728 | - | 1 | 1% | 47% | 21% | 409 |
| Amazon 14 | Gaillardet et al. 1997 | Amazon | Wester Tributaries & Amazon Mainstem | 0.71112 | - | 1 | 1% | 47% | 21% | 409 |
| Amazon 20 | Gaillardet et al. 1997 | Amazon | Wester Tributaries & Amazon Mainstem | 0.711478 | - | 1 | 2% | 42% | 25% | 429 |
| Amazon 6 | Gaillardet et al. 1997 | Amazon | Wester Tributaries & Amazon Mainstem | 0.709172 | - | 1 | 0% | 60% | 17% | 407 |
| Rio Madeira | Gaillardet et al. 1997 | Lower Madeira | Beni-Madeira & Lower Negro | 0.720036 | - | 1 | 2% | 20% | 30% | 412 |
| Rio Negro | Gaillardet et al. 1997 | Lower Negro | Beni-Madeira & Lower Negro | 0.716223 | - | 1 | 0% | 21% | 68% | 1243 |
| Rio Topajos | Gaillardet et al. 1997 | Lower Topajos | Lower Amazon Tributaries | 0.733172 | - | 1 | 10% | 8% | 48% | 658 |
| Rio Trombetas | Gaillardet et al. 1997 | Lower Trombetas | Lower Amazon Tributaries | 0.732295 | - | 1 | 3% | 7% | 78% | 515 |
| Uracara | Gaillardet et al. 1997 | Lower Uracara | Lower Amazon Tributaries | 0.723584 | - | 1 | 2% | 19% | 62% | 558 |
| Purus | Queiroz et al. 2009 | Lower Purus | Wester Tributaries & Amazon Mainstem | 0.711135 | - | 1 | 0% | 92% | 4% | 564 |
| Solimões* | Queiroz et al. 2009 | Lower Solimões | Wester Tributaries & Amazon Mainstem | 0.714461 | - | 1 | 0% | 63% | 1% | 289 |
| Atalaya | Santos et al. 2013 | Upper Solimões | Wester Tributaries & Amazon Mainstem | 0.70887 | - | 1 | 0% | 20% | 0% | 331 |
| Borba | Santos et al. 2013 | Lower Madeira | Beni-Madeira & Lower Negro | 0.71762 | - | 1 | 2% | 20% | 30% | 412 |
| Borja | Santos et al. 2013 | Upper Solimões | Wester Tributaries & Amazon Mainstem | 0.7085 | - | 1 | 0% | 13% | 0% | 426 |
| Caracarai | Santos et al. 2013 | Upper Negro | Beni-Madeira & Lower Negro | 0.72238 | - | 1 | 0% | 0% | 74% | 1650 |
| Francisco de Orellana | Santos et al. 2013 | Upper Solimões | Wester Tributaries & Amazon Mainstem | 0.70592 | 0.00037 | 26 | 0% | 67% | 10% | 287 |
| Itiatuba | Santos et al. 2013 | Lower Tapajos | Lower Amazon Tributaries | 0.72964 | 0.00587 | 27 | 10% | 4% | 51% | 671 |
| LaBrea | Santos et al. 2013 | Lower Purus | Wester Tributaries & Amazon Mainstem | 0.71012 | - | 1 | 0% | 90% | 6% | 1126 |
| Manacapuru | Santos et al. 2013 | Lower Solimões | Wester Tributaries & Amazon Mainstem | 0.70907 | 0.00025 | 38 | 0% | 72% | 2% | 312 |
| Obidos | Santos et al. 2013 | Amazon | Wester Tributaries & Amazon Mainstem | 0.71154 | 0.00053 | 46 | 1% | 46% | 23% | 408 |
| PortoVelho | Santos et al. 2013 | Lower Madeira | Beni-Madeira & Lower Negro | 0.71677 | 0.00073 | 9 | 3% | 17% | 22% | 379 |
| Ruranbaque | Santos et al. 2013 | Upper Madeira | Beni-Madeira & Lower Negro | 0.71730 | 0.00126 | 38 | 17% | 13% | 1% | 433 |
| Serrinha | Santos et al. 2013 | Upper Negro | Lower Amazon Tributaries | 0.73183 | 0.00737 | 16 | 0% | 19% | 80% | 614 |
| Tabitinga | Santos et al. 2013 | Upper Solimões | Wester Tributaries & Amazon Mainstem | 0.70881 | 0.00029 | 9 | 0% | 75% | 3% | 200 |
| Amazon Mouth (Predicted) | Predicted from regression | Amazon | Wester Tributaries & Amazon Mainstem | 0.71625 | 0.00787° | - | 2% | 40% | 29% | 482 |
| Jari (Predicted) | Predicted from regression | Lower Jari | Lower Amazon Tributaries | 0.72928 | 0.00873° | - | 0% | 2% | 89% | 718 |
| Paru (Predicted) | Predicted from regression | Lower Paru | Lower Amazon Tributaries | 0.72703 | 0.00845° | - | 1% | 4% | 78% | 515 |
| Tocantins (Predicted) | Predicted from regression | Lower Tocantins | Lower Amazon Tributaries | 0.72683 | 0.00884° | - | 13% | 22% | 36% | 877 |
| Xingu (Predicted) | Predicted from regression | Lower Xingu | Lower Amazon Tributaries | 0.72633 | 0.00827° | - | 2% | 13% | 70% | 1091 |

\* Outlier dropped from regression analysis
° Values are prediction intervals of the regression from Equation 1.

**Figure 3.**
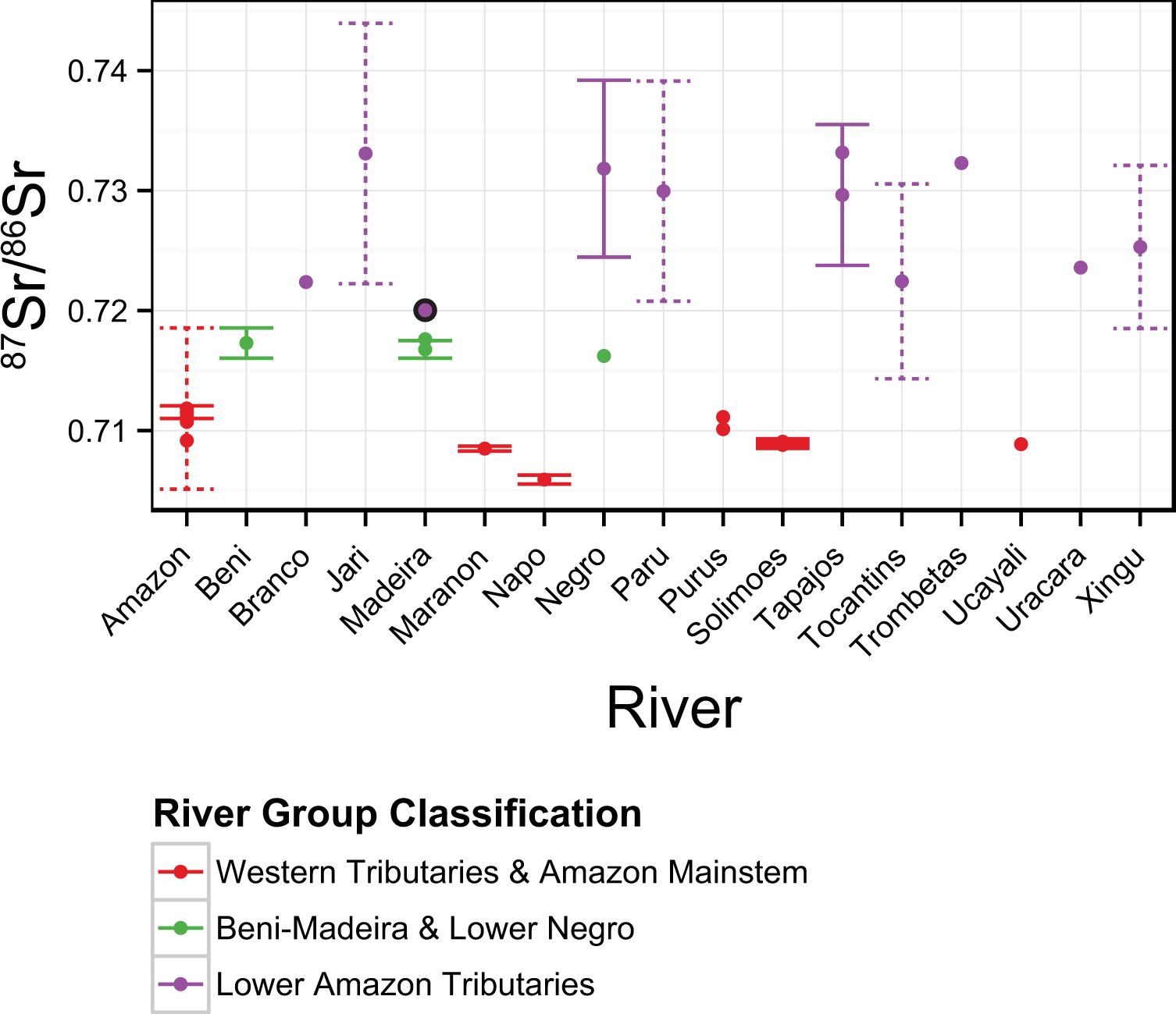
River isotopic signatures throughout the Amazon River basin. Strontium ratio values (y-axis) for each sampled and predicted watershed (x-axis) in the current study. Color indicates the classification to three river groups using quadratic discriminant function analysis. Solid error bars indicate the standard deviation where samples were repeated over time (See Table 1 for sample sizes). Dashed error bars indicate the prediction intervals from the geologic regression (Equation 1) used to predict that point. Points bordered in black were misclassified during cross validation of the quadratic discriminant function.

The isotopic chemistry of a few significant locations were not available in the literature. Notably missing were samples from the mouth of the Amazon River, its tributaries below Obidos, and the Tocantins River which contributes to the estuary habitat of our study species. To account for the ^87^Sr/^86^Sr of these locations we used the relationship between the geologic makeup of a watershed and its ^87^Sr/^86^Sr signature to predict these points, following a similar regression approach to Hegg et al. [54]. Watersheds were delineated in qGIS (http://www.qgis.org), an open-source geographic information system, using the GRASS analysis plugin, which contains advanced watershed analysis tools from the open-source GRASS GIS platform (http://grass.osgeo.org). All analysis layers were procured from open-access datasets. Water sampling points were manually digitized by the authors based on location descriptions from Santos et al. [53], Gaillardet et al. [52], and Queiroz et al. [51]. Topography layers were taken from the GTOPO30 dataset [59] and stream courses from the HydroSHED dataset [60]. Geologic information came from the World Energy Assessment Geologic Map of the Amazon Region [61].

We used geologic age as the primary candidate independent variables in our regression to predict the ^87^Sr/^86^Sr signatures for unsampled tributaries, along with very general intrusive and extrusive rock-type categories (Figure 1B). Our methods differed from Hegg et al. [54], who used rock type as the primary explanatory variable rather than age. We did this because the very generic designations of intrusive or extrusive rock available in our dataset were insufficient to provide explanatory power. The values for these candidate variables were calculated by converting the geologic age codes from the map attribute table to continuous variables using the mean age (Ma) of the geologic periods encompassed by each code using the International Chronostratigraphic Chart [62]. The percentage area of each rock age and type was then calculated within the watershed upstream of each ^87^Sr/^86^Sr sample point. The mean age of each watershed, weighted by area, was also included as a potential explanatory variable for the regression, leaving twenty-four potential explanatory variables for the regression.

Model selection used a genetic algorithm selection procedure in the {glmulti} package for R [63]. We limited models to four terms to limit the number of potential models and included interaction terms. The genetic algorithm uses a search algorithm based on Darwinian natural selection, an efficient method for model optimization when the number of potential models is large, as was the case with our geologic data [64]. Akaike’s Information Criterion optimized for small datasets (AICc) was used as the optimization criteria for the genetic algorithm, a criterion that penalizes over-parameterization [65]. One third of the sample points were randomly selected as a validation set, withheld from the model selection procedure, and used to assess prediction accuracy of the best model. The best model was then used to calculate the ^87^Sr/^86^Sr values for the unsampled points in the basin, using the geology upstream of these points.

### Grouping of Distinguishable Watersheds

The water sample points were grouped into three distinguishable geographic regions using prior knowledge of the geography and geology of the watersheds (Table 1, Fig. 1A, Fig. 3). River basins that were geographically contiguous and broadly geologically and chemically similar were grouped. The Amazon River mainstem and western tributaries, all considered whitewater rivers [26], were grouped together due to the overwhelming influence of the Andes on their chemistry. The Beni-Madeira River and lower Negro River were grouped due to similar chemical signatures from a mix of upland mountainous geology and old, lowland, Amazon and Guyana shield geology. The Negro River, being blackwater, would not be expected to contain large numbers of our target species, while the whitewater Madeira is a known fishery [23,26]. The lower Amazon tributaries (below the Madeira River) were grouped due to the their similarly old, shield geologies resulting in high ^87^Sr/^86^Sr values. These rivers are all considered clearwater tributaries [26]. These group assignments were then used as the training set, with ^87^Sr/^86^Sr as the predictor and group as the response, to create a quadratic discriminant function. This discriminant function was then used in the following section to classify the ^87^Sr/^86^Sr signatures recovered from fish otoliths to these three distinguishable river groups.

### Determining Fish Movement and Location

The transect of ^87^Sr/^86^Sr values from the core to the rim of each otolith was analyzed using a PELT algorithm changepoint analysis (using the {changepoint} package in R [66]) to determine when the mean ^87^Sr/^86^Sr values changed to a new stable signature. This changepoint algorithm generated mean values and starting points for each stable region within the otolith transect using a penalty value of 0.0001 [67]. Each stable signature was assumed to correspond to movement into a new river signature, with the first stable signature corresponding to early rearing. In some cases the changepoint algorithm returned erroneous means for small portions of the signature, in locations were the means was obviously unstable. Such fragments were manually removed.

Stable otolith signatures were then classified to their likely location using the discriminant function developed from known and predicted water sampling points in the prior section. Because *a priori* probability of group membership was unknown, the prior probabilities for the discriminant function were set to be equal among groups. After classification, the results were plotted and the data were assessed to determine trends in early rearing and movement both within and among the three sampled species.

## Results

### Baseline Water Sampling and Prediction

The most parsimonious model without interaction terms explained ∼80% of the variation in the data but provided an unreasonably high prediction for the mouth of the Amazon River. We had no direct evidence of significant geologic interactions; however, we included interactions in a second model selection exercise in hopes of finding a parsimonious model that better fit the available data. We limited the maximum number of model terms to four to limit the number of potential models available from the twenty-four available variables plus interactions. This limit is reasonable since more terms would risk over parameterization given the number of observations used to build the model. Under these conditions the AICc model-selection algorithm selected three models that were greater than two AICc points superior to the next most parsimonious model. The top model,

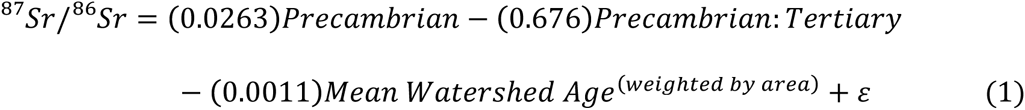

explained 89% (Adjusted R^2^) of the variation in the data, provided the best prediction residuals for the validation set, and resulted in a more reasonable prediction of the mouth of the Amazon. This equation was used to predict the ^87^Sr/^86^Sr signatures for the five unsampled watersheds.

### Grouping of Distinguishable Watersheds

A quadratic discriminant function provided the best cross-validation error rate (3.6%) for discriminating all the watersheds into the three regions. One predicted value for the Madeira River was the lone misclassification from the validation set, being classified to the Lower Amazon Tributaries group. One sample from the Solimões River was dropped from the training set as an outlier (Table 2). The value for this site was unexplainably high in comparison to the multi-year samples above and below it on the river. While no explanation for this discrepancy was forthcoming from the original study, Bouchez et al. [68] found that lateral heterogeneity in ^87^Sr/^86^Sr signatures can persist for long distances below confluences in the Solimões.

### Determining Fish Movement and Location

Changes in ^87^Sr/^86^Sr ratio, indicating movement, were common across each of the three species. Movement between distinguishable river groups, as determined by discriminant function classification, was less frequent. Over 70% of dourada exhibited movement between stable signatures based on changepoint analysis, however only two (14%) showed movement between distinguishable river groups after discrimant function classification (Figure 4A). Sample BR24 started life in the Amazon Mainstem and Western Tributaries signature before moving to a signature consistent with the Lower Amazon Tributaries river group. Sample BR25 began life with a signature consistent with the Lower Amazon Tributaries river group, before moving twice to a signature consistent with the Beni-Madeira and Lower Negro group with a small region consistent with the Amazon Mainstem and Western Tributaries river group.

**Figure 4.**
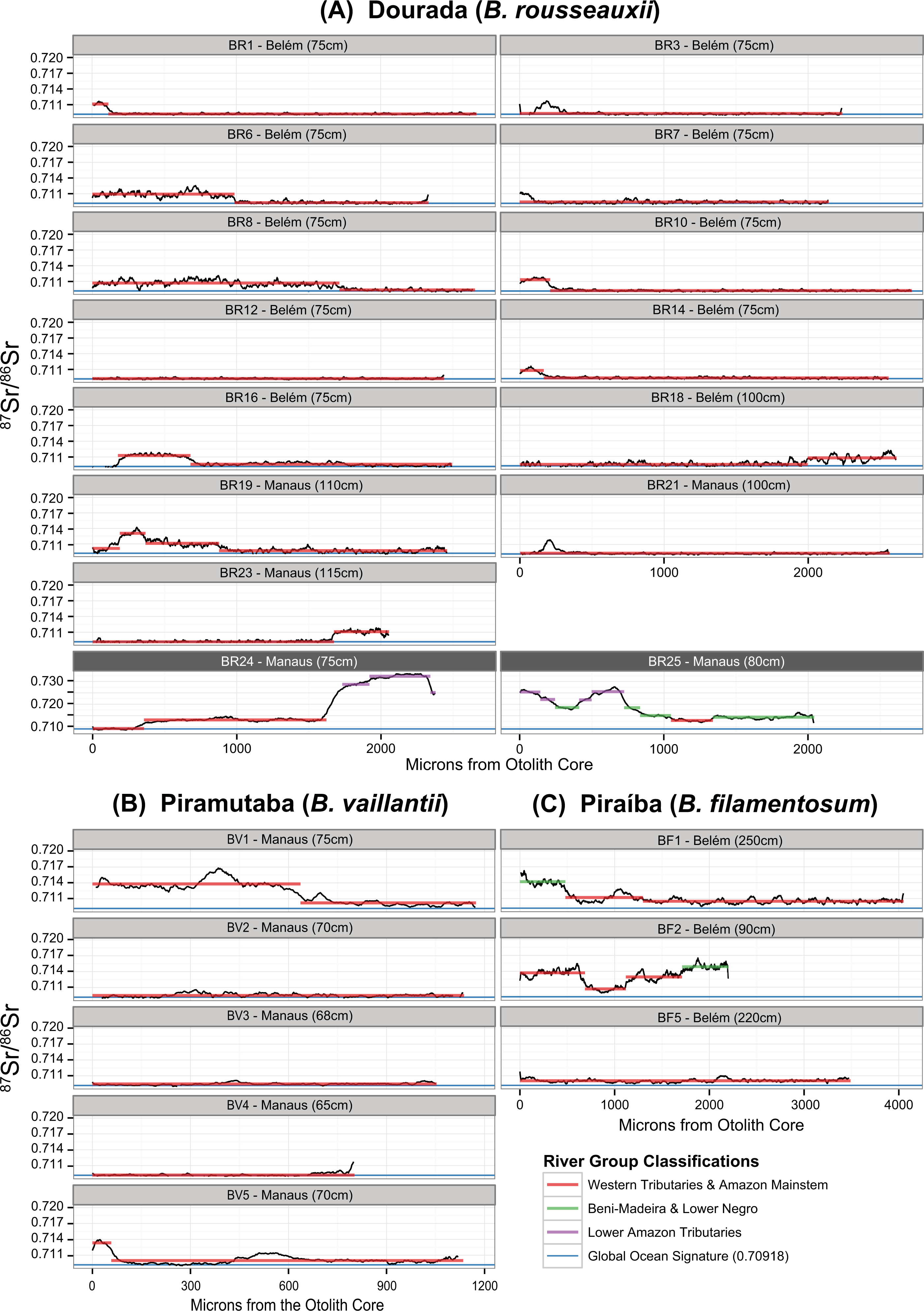
Location classification of ^87^Sr/^86^Sr signatures in otolith transects. Plots show the variation in ^87^Sr/^86^Sr (y-axis) over the life of sampled fish, represented as distance from the otolith core in microns (x-axis). Horizontal colored lines indicate stable signatures identified using changepoint analysis, with colors representing inclusion in one of three statistically distinguishable river groups based on quadratic discriminant analysis. Fourteen samples of dourada (A) were collected in Belém and Manaus fish markets. Five samples of piramutaba (B) were collected in Manaus. Three samples of piraíba (C) were collected in Belém. Dark grey chart labels indicate a different y-axis scale was used to accommodate large variations in ^87^Sr/^86^Sr. The x-axis scale differs for all fish depending on the width of the otolith, which varies based on age, growth and species specific factors.

Of five piramutaba, two (40%) were shown to move between stable signatures but none moved outside the Amazon Mainstem and Western Tributaries river group based on discriminant function classification (Figure 4B). Of the three piraíba samples two showed movement, both of which moved between the Beni-Madeira and Lower Negro River group and the Amazon Mainstem and Western Tributaries (Figure 4C).

Overall, these results indicated that the majority of fish begin life in the Amazon Mainstem and Western Tributaries signature, a signature which contains within it the estuary signature, the expected location of early rearing [23]. Our results were unable to distinguish the Amazon mainstem signature from that of the estuary, the expected location of early rearing. However, we would expect the signature to be intermediate between that of the signature of the Amazon River mouth and the global marine signature of 0.70918 resulting from the mixing of the two water bodies. The resulting estuary signature would be contained with the range of the Amazon Mainstem and Western Tributaries classification.

Conversely, some portion of fish begin life, and undergo some or all of their rearing, in signatures indicative of freshwater (Figure 4A, B &C). Sample BR25 (dourada) and BF1 (piraíba) were the two most obvious examples, starting life with signatures that correspond to the Beni-Madeira and Lower Negro and Lower Amazon Tributary signatures. Many of the fish that remained within the Amazon Mainstem and Western Tributaries river group began life with signatures > 0.71, significantly higher than the Mainstem Amazon River signatures (Table 2)and the global marine signature, and thus likely an upriver signature. Others (BR6, BR8, BV1, BF2) spent large portions of their lives in distinctly upriver environments.

Otolith microchemistry data is available in the supporting information for this publication (S1 Table).

## Discussion

Understanding the migration ecology of Amazon catfish represents an opportunity to understand the large-scale ecosystem processes of long distance aquatic migration in an intact, native fish population. These fish also represent an international conservation challenge, as their movements stretch across multiple national boundaries [35] and are potentially threatened by several hydropower projects [16–19]. The first step in understanding these ecological processes, and their effects on the fishery, is a robust understanding of their migratory movements at a finer scale than is currently available from fisheries data.

Our results provide the first reported reconstruction of movements and migrations of individual *Brachyplatystoma* spp. in the Amazon Basin using chemical signatures from otoliths. Applying these techniques in this system appears promising for improving our understanding of the migratory movements of these species, which are currently understood only at the most basic level. While this work is preliminary, ^87^Sr/^86^Sr signatures appear capable of identifying large-scale fish movements at a meaningful spatial scale; between important river systems within the basin. Further, movement between areas with stable signatures can be observed within statistically distinguishable river groups, indicating that improved baselines and larger sample collections should allow greater resolution. Discrimination of fish location and movement may also be improved by applying multivariate analyses of elemental signatures in concert with ^87^Sr/^86^Sr signatures [69–71].

We observed diversity in rearing location and behavior among a minority of individuals in each of the three species sampled using otolith ^87^Sr/^86^Sr movement reconstructions and discriminant function classification to location. For example, fish BR25 (a dourada caught in Manaus) appears to have spent most of its early life in areas with very high ^87^Sr/^86^Sr ratios assigned to the Lower Amazon Tributaries river group (Figure 4A). The chemical signatures of this fish indicate that rather than drifting to the estuary to rear, it spent the first third of its life in a lower tributary of the Amazon River before moving to a signature indicative of areas between the Madeira and Amazon Rivers, finally being caught near between Manaus and the mouth of the Madeira (as reported by the fishermen who provided the samples in Manaus). Notably, the signatures appear to support the idea that this fish never traveled to the estuary. Fish BF1 (piraíba) also showed a significant length of freshwater rearing in a signature assigned to the Beni-Madeira and Lower Negro river group (Figure 4C).

Indeed, several additional dourada and piraíba, and one piramutaba all appeared to have spent significant time in ^87^Sr/^86^Sr signatures greater than 0.7100 (Figure 4A, B &C). This signature is higher than all but one location on the mainstem Amazon River and significantly higher than the accepted global marine signature of 0.70918 considering the analytical precision that is possible for otolith measurements. We would expect estuarian signatures to fall between the ^87^Sr/^86^Sr signature of the Amazon mouth and the global marine signature except very near the mouth of the Tocantins. The Tocantins signature is likely attenuated significantly by Amazon River water flowing through the Canal de Breves which noticeably muddies the water flowing from the Tocantins [23]. Further, the estuary signature should converge to the marine signature at relatively low salinities due to the much higher strontium concentration of the ocean [72]. This indicates the possibility that a larger percentage of our samples may have reared upriver of the estuary as well, but our methods were unable to detect it.

Ecologically, the presence of upriver ^87^Sr/^86^Sr signatures during the rearing phase indicates that in some situations a rearing strategy that forgoes the high growth potential of the estuary may provide overall fitness benefits. This finding suggests that the life-history of these species is more complex than has been previously understood. The potential existence of a freshwater rearing life-history, based on evidence for alternative spawning periods in the upper reaches which do not fit the conventional estuary rearing model, was theorized by García Vasquez et al. [33]. This model is supported by evidence of young and immature fish present in the far western Amazon before and during the spawning season when the prevailing hypothesis would place these fish in the estuary [24,35,36].

Understanding the extent of diversity in life history strategies is critical for managing these little-studied native species, especially in habitats facing current and future perturbations. Variations in life history have been shown to affect recruitment, survival, and fisheries sustainability in other long-distance migratory fish [73–75]. These species may home to their natal rivers, and sub-population structure and associated differences in life history may exist [37,38], which may have important implications for planning and policy decisions related to dam placement and fishery management. For instance, fishing pressure in the estuary is high, potentially limiting juvenile escapement to upriver fisheries [24,35]. Freshwater rearing life-histories would avoid the high fishing pressures in the estuary, potentially increasing survival and providing important recruitment to upriver fisheries in the Western Amazon, fisheries which appear to be overfished [24,25,31].

Dams in particular have been shown to decrease life-history diversity of other major migratory fisher species, with consequences for their conservation and fisheries sustainability [76,77] and diverse source populations appear to provide resilient fisheries over time [74]. So, the extent to which dams or other anthropogenic disturbances decrease the diversity of source populations in dourada, piramutaba, and piraíba could have adverse impacts on the sustainability of the fishery. The pace of dam building in the major Amazon Basin tributaries [16,17,19]; the difficulty of providing significant fish passage in Amazonian rivers [21]; and the known and suspected effects of dams on the migration of these migratory catfish species [6,29,78] increase the need to understand the details of their migration ecology. Only with detailed knowledge of *Brachyplatystoma* migration ecology can policymakers weigh the effects of dams on the sustainability of this important fishery.

Our study raises numerous important questions and opportunities for future research. While it is clear that water chemistry signatures for the Amazon basin can be classified into meaningful groups, such results do not necessarily translate into concrete interpretation of fish movement at anything but the largest of scales. Our understanding of the degree to which the movements and migrations of individual fish can be interpreted at the scale of the entire Amazon basin is in its infancy. At smaller spatial scales, many of the local tributaries to the major rivers of the Amazon basin exhibit vastly different signatures than those of the main channels, which reflect the headwater signature of the Andes or the Brazilian and Guyana Shield geology [51]. Incomplete or slow mixing of different signatures, especially across muddy whitewater and tannic blackwater rivers, may occur over extended river distances, which may create unexpected intermediate signatures [68]. Furthermore, our understanding of temporal variation associated with available empirical data is limited. Especially in the Amazon estuary our understanding of the ^87^Sr/^86^Sr signatures in this seasonally dynamic environment are limited. Increased understanding of these estuarian signatures is critically important to understanding these species using otolith microchemistry studies. Our study species are thought to inhabit only the larger, mainstem tributaries, making large scale location classification useful. At smaller geographic scales additional ground truth sampling of fish and water chemistry is needed to constrain the scale at which movements and migrations can be accurately interpreted from otolith data. Otolith sampling across larger areas of the Amazon may also allow elemental ratios, which are fractionated biologically unlike ^87^Sr/^86^Sr, to be used in addition to ^87^Sr/^86^Sr to improve location analysis.

Overall, this study highlights the feasibility and utility of the latest otolith chemistry techniques to greatly improve our understanding of the movements and ecology of these important native fishes throughout the entire Amazon basin. Recent declines in the fishery point to the necessity of conducting this research [22,28,79]. Migratory catfish in the Amazon Basin are several of only a few large, freshwater fishes worldwide that are not currently imperiled due to anthropogenic changes to freshwater ecosystems [12]. However, as fishing pressure increases, land use and forest clearing affect the river system, and dams threaten migration routes and access to critical habitats, these populations will likely be affected. The sustainability of these populations and the fisheries they support, especially across international borders, continues to depend on accurate population assessments of based on detailed knowledge of their behavior and ecology.

## Acknowledgements

Thanks to A. Fremier and P. Anders for their encouragement and help in framing the project at the earliest stages, as well as sharp outside editing. Thanks to C. Cooper for editing and moral support. Thanks to L. Rayala for applying his practiced editing eye. Thank you to Z. Hogan for providing constructive feedback on the project proposal. Thanks to M. Andrade, A. Zuluaga and D. Bastos for help with sample collection in Manaus and Belém, as well as to the fishermen who volunteered their catch for sampling. Thanks to J. Vervoort and C. Knaack at the Washington State University Geoanalytical Laboratory for use of their equipment for isotopic analysis. T. Giarrizzo received a productivity grant from CNPq (process: 308278/2012-7), and was funded by CAPES (PNPD and Pró-Amazônia: biodiversidade e sustentabilidade).

## Supporting Information

**S1 Table. Otolith transect data.** Table of raw otolith transect data for each sample analyzed.

